# Damsel: Analysis and visualisation of DamID sequencing in R

**DOI:** 10.1101/2024.06.12.598588

**Authors:** Caitlin G Page, Andrew Londsdale, Katrina A Mitchell, Jan Schröder, Kieran F. Harvey, Alicia Oshlack

## Abstract

**Summary:** DamID sequencing is a technique to map the genome-wide interaction of a protein with DNA. Damsel is the first Bioconductor package to provide an end to end analysis for DamID sequencing data within R. Damsel performs quantification and testing of significant binding sites along with exploratory and visual analysis. Damsel produces results consistent with previous analysis approaches.

**Availability and implementation:** The R package Damsel is available for install through the Bioconductor project https://bioconductor.org/packages/release/bioc/html/Damsel.html and the code is available on GitHub https://github.com/Oshlack/Damsel/

**Supplementary information:** Available through the journal.

## INTRODUCTION

Defining the binding landscape of chromatin interacting proteins is vital for understanding and manipulating gene expression. DamID was developed as an alternative to ChIP-seq that facilitates this without relying on antibodies (van Steensel and Henikoff, 2000). In the DamID protocol, the E.coli DNA adenine methylase (Dam) protein is fused to a transcription regulatory protein of interest, and upon interaction with DNA, Dam methylates the adenine in adjacent GATC motifs. As Dam can also randomly methylate GATC motifs, DamID analysis is always conducted with the expression of a Dam-only control. Dam-methylated genomic regions are enriched using a methylation sensitive endonuclease to cut DNA and amplified with PCR. The ends of the resultant DNA fragments are sequenced to provide the genome-wide binding profile of the protein of interest. Following the identification of enriched genomic regions relative to the Dam-only control, potential target genes of the protein of interest can be identified. These target genes are identified based on their proximity to a peak. A limitation of DamID is the reliance on GATC motifs, which limits the utility of DamID to genetically tractable species. Targeted DamID is an optimised version of DamID that was developed in Drosophila melanogaster which allows in vivo expression and precise spatiotemporal control (Southall et al., 2013, Marshall et al., 2016, Aughey et al., 2019).

Despite DamID being a powerful alternative to ChIP-seq, there is limited availability of analytical tools. Bespoke analyses are commonly conducted, as in Vissers et al. (2018) which utilises edgeR’s (Robinson et al., 2010) robust statistical testing. A 2019 review of DamID mentions only two published methods, both focussed on algorithms to facilitate peak calling (identifying regions of binding compared with the control) (Aughey et al., 2019). Neither of these tools conduct the whole DamID bioinformatics workflow, thereby forcing researchers to rely on combining multiple tools and platforms with bespoke analysis. The widely used method, damidseq_pipeline (Marshall and Brand, 2015), is the most comprehensive tool and runs on the command line analysing samples individually, but has no visualisation capabilities. Here, we present Damsel, the first dedicated R package for DamID, providing an end-to-end analysis, including exploratory and visualisation capabilities.

## METHOD

The Damsel workflow, as described in Figure 1A, starts after read alignment using standard tools (Supplemental Methods) and requires BAM files and GATC regions for a genome as input. Damsel conducts the following steps in the analysis workflow: region quantification, identification of bound genomic regions, peak calling, candidate gene detection, and gene ontology testing, described in more detail below.

**Figure 1.**
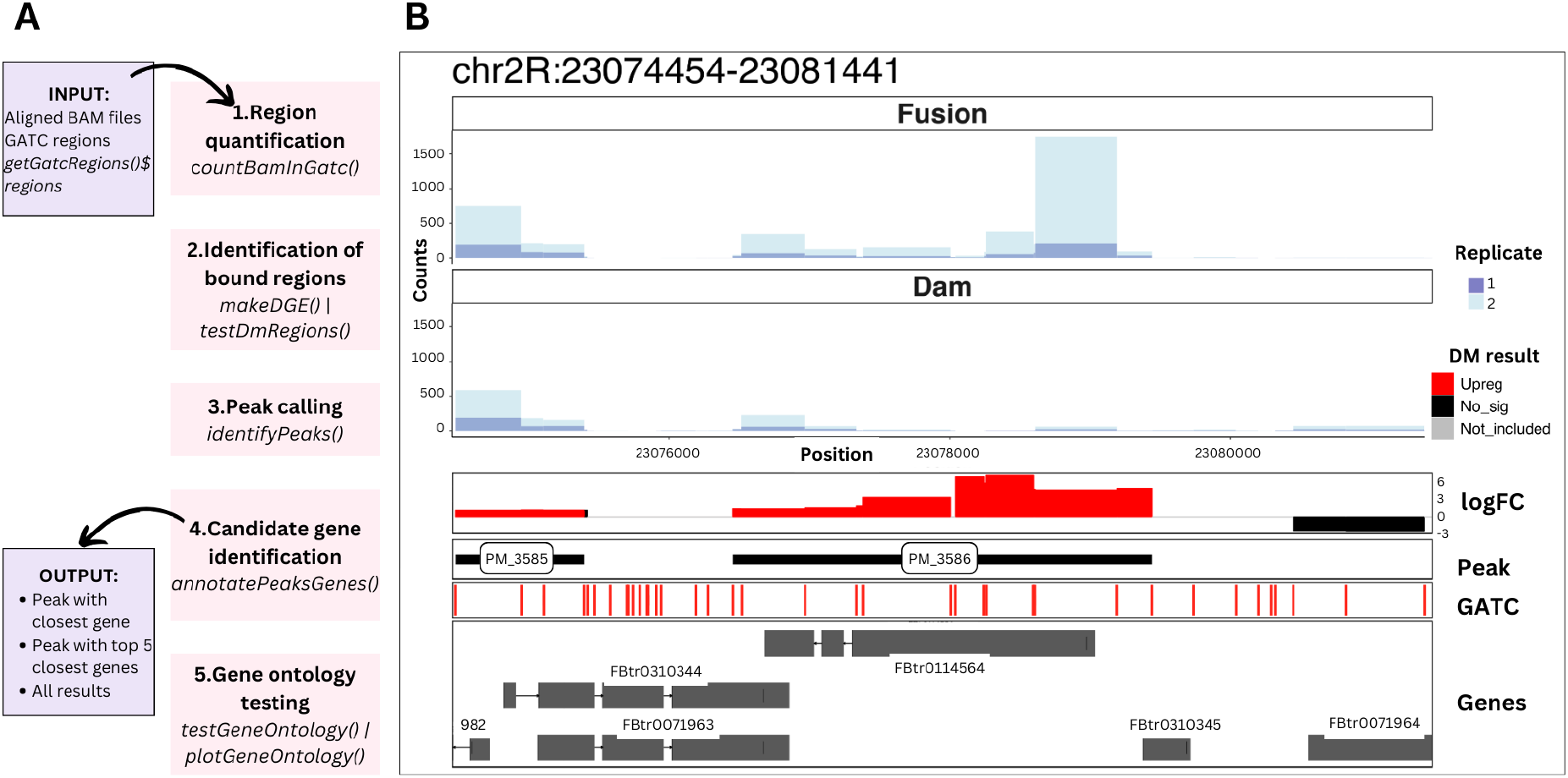
Overview of Damsel. **A** The main steps in the Damsel workflow alongside their Bioconductor functions, including region quantification, identification of bound regions, peak calling, candidate gene detection, and gene ontology testing. **B** Example visualisation available in Damsel of the most significant peak found within the Vissers data. The layers of the plot show the counts across the regions for the Fusion and Dam-only samples, the differential methylation test results, the presence of significant peaks, the positions of the GATC sites, and the overlap to gene annotations.

### 1. Region quantification

The genome is intrinsically partitioned into genomic regions demarcated by GATC motifs (note that GATC is its own reverse complement and therefore the motif demarcates both strands of the DNA simultaneously). Damsel then utilises the *featureCount()* function from the Rsubread package (Liao et al., 2019) to summarise the read counts between two adjacent GATC sites (GATC region) from the provided BAM files and motif input file. For paired-end BAM files, Damsel instead summarises the fragments (read pairs). Due to the background methylation that occurs in the Dam-only sample, there is high correlation of the read counts of the regions between samples (Supplementary Fig. S1).

### 2. Identification of bound genomic regions

Previously, enrichment of a bound genomic region was identified from the log2 ratio of the region counts of the fusion sample compared to the control. However, analysis by Marshall and Brand (2015) established that this can result in a negative bias, highlighting the importance of normalisation. Damsel first filters out excessively large regions (> 10 000 bp), along with low count regions, before normalisation and statistical testing is conducted with edgeR, incorporating the replicates for a streamlined analysis (Robinson et al., 2010). edgeR’s TMM normalisation assumes the majority of regions are not significant (Robinson and Oshlack 2010), and differential testing utilises a quasi-likelihood negative binomial model (Robinson et al., 2010), utilising experimental replicates to model the biological variance in the data.

#### 3. Peak calling

Peak calling allows for aggregation of adjacent significant regions that are likely to be the result of the same binding event. Damsel identifies peaks by aggregating the region level differential expression results, with p-value threshold set to 0.01 (by default), and log2 fold change threshold set to one. If the gap to the next significant region is less than 150bp, it is included in the peak. Regions smaller than this size have consistently lower counts, and thus represent a greater proportion of regions that are excluded from differential testing. Peaks are ranked based on the statistics for ChIP-seq analysis presented in csaw (Lun and Smyth 2016), using the region with the lowest p-value to represent the peak and ordering peak significance based on this p-value.

### 4. Candidate gene identification

Methods for associating peaks and genes match the peak to its closest gene, with distance taken from the middle of the peak to the transcription start site. This is straightforward if the analysis is conducted on a species with little overlap between genes, like humans. However in species like *Drosophila melanogaster*, there is a large amount of overlap between genes and the closest gene may not be the most likely candidate. Therefore, Damsel outputs each peak with a list of candidate target genes, along with their genomic positions and relevant statistics. Placing the genes within the context of the peaks provides a more informative output, and sets Damsel apart from other tools.

At this point, Damsel outputs a file with all identified peaks and their associated statistics, matched to their associated candidate genes. Comparatively, damidseq_pipeline (Marshall and Brand 2015) outputs a list of genes and a score, which is the end of the analysis. However, Damsel also includes the following utilities.

### 5. Gene ontology testing

Gene ontology testing can be used to interpret the set of candidate target genes identified from the peaks. Damsel repurposes goseq, an R package developed to correct for detection biases in RNA-seq (Young et al., 2010). This is important because the GATC motifs are not spread evenly throughout the genome and different genes have different lengths and therefore a different number of GATC regions associated with them. This results in a bias where genes with more GATC regions are more likely to be associated with a peak (binding event) (Supplementary Fig. S2). Without bias correction, the gene ontology terms identified are not truly representative of the data. goseq takes as input a list of genes, if they are significant, and the number of GATC regions within 2kb of the gene as the bias parameter, and outputs a list of overrepresented gene ontology terms.

### 7. Visualisations

A unique feature of Damsel is its visualisation capability. Other DamID methods, including damidseq_pipeline (Marshall and Brand 2015), output files that they recommend viewing in IGV, requiring researchers to switch software platforms. Damsel provides the opportunity to use its output files to create plots within R. Building on ggbio (Yin et al., 2012) and ggcoverage (Song and Yang 2023), Damsel contains ggplot2 (Wickham 2010) style plots that present results from different stages of the analysis. These plots can be layered, allowing the visualisation of counts per region, log2 fold change and differential expression testing p-values, peak location, GATC motif, and gene positions for a provided region (Figure 1.B).

## RESULTS

The existing approaches for DamID analysis (Marshall and Brand (2015), Vissers et al., (2018)) conduct a similar sequence of stages as Damsel including: identifying regions of enrichment, combining regions into peaks, and associating peaks with genes (Supplementary Table 1). Some of these stages are handled differently in Damsel with increased functionality and seamless data flow through the pipeline. Like Vissers’ (2018) analysis, Damsel incorporates replicates into the analysis to model variability and significantly reduce the burden of compiling results. Additional features unique to Damsel include region quantification directly from bam files, the ranking of significant results (peaks and genes), providing relevant statistics associating the peaks with annotated genes, and capacity for visualisation of the results (Supplementary Table 1).

To evaluate Damsel, we applied it to a subset of the data presented in Vissers et. al. (2018). To evaluate the false positive rate, the samples from this experiment were randomly assigned as Fusion and Dam-only. When a Dam-only and Fusion sample from different replicates were assigned as Dam-only (or Fusion), the results were as expected - with no regions identified as significant at FDR < 0.01, and therefore no peaks were identified. This demonstrates good false discovery rate control. Similarly the Vissers’ pipeline and the Marshall and Brand pipeline (2015) found no significant peaks in this scenario.

We next analysed the data in the way it was intended with two replicates of the fusion and control. Damsel was found to be more sensitive than the original reported results finding 2,955 peaks, which is 669 more than identified from running Vissers’ (2018) pipeline. Unsurprisingly, Damsels results were more similar to the Visser pipeline than to DamID-seq pipeline with more than 60% of peaks overlapping with Visser (Supplementary Figure S3). We performed Damsel’s GO testing on the peaks reported in the original paper and those found by Damsel and reassuringly found that Damsel reported 9 of the same top 10 GO ontology terms as the regions from Vissers (Supplementary Figure S4). Finally, as an example of data visualisation we used Damsel to show the data and analysis of the most significant peak identified by all three methods (Figure 1B).

Identifying which method is the most sensitive and accurate is beyond the scope of this study, and would be difficult without a large amount of experimental validation. However, Damsel performs consistently compared with established methods, and has produced results on multiple published datasets.

## CONCLUSION

Damsel offers an end-to-end DamID analysis pipeline including exploratory analysis and visualisations. On published data we find similar results to previous bespoke analysis. We find that Damsel can control false discovery rate and is more sensitive. Damsel includes new features such as the ranking of peak significance and performing gene ontology analysis. Future work will include the utility to perform motif enrichment analysis on sequences found in enriched peaks.

## Data availability

The data used in this paper is available at the European Nucleotide Archive (PRJNA494322) https://www.ebi.ac.uk/ena/browser/view/PRJNA494322

## Acknowledgements

We thank Avni Avand for testing the software and providing feedback.

## Funding

A.O was supported by an Investigator grant (APP1196256) from the National Health and Medical Research Council of Australia. K.F.H was supported by an Investigator grant (APP1194467) from the National Health and Medical Research Council of Australia. *Conflict of Interest:* none declared

## Notes

https://www.ebi.ac.uk/ena/browser/view/PRJNA494322

### Competing Interest Statement

The authors have declared no competing interest.

