## Supplementary Information for "Damsel: Analysis and visualisation of DamID sequencing in R"

**SUPPLEMENTARY METHODS:**

1. Processing data

The example data used in this paper is a subset of the data created by Vissers' et al., (GSE120731, SRA:SRP163124). The following samples were used in the subsequent analysis:

- GSM3409439 Dam-only Rep 1
- GSM3409440 Dam-Sd Rep 1
- GSM3409441 Dam-only Rep 2
- GSM3409442 Dam-Sd Rep 2

FastQ files were downloaded from the European Nucleotide Archive (https://www.ebi.ac.uk/ena/browser/view/PRJNA494322) with separate fastq files for the paired end samples Dam-only Rep 1 and Dam-Sd Rep 1.

FastQ files were aligned using RSubread. The drosophila reference genome (BDGP6.46) was obtained from EnsemblMetazoa (http://ftp.ensemblgenomes.org/pub/metazoa/release-59/fasta/drosophila_melanogaster/dna/).

```
buildindex(basename = "dros_ref", reference =
"path/Drosophila_melanogaster.BDGP6.32.dna.toplevel.fa")
```

- For the single end fastq files (dam-only rep 2, dam-sd rep 2)

```
align(index = "dros_ref", readfile1="path/SRR7948877.fastq")
```

- For the paired end fastq files (dam-only rep 1, dam-sd rep 1)

```
align(index = "dros_ref", readfile1 = "path/SRR7948872_1.fastq.gz",
readfile2 = "path/SRR7948872_2.fastq.gz")
```

The subsequent BAM files were then sorted and indexed using samtools.

```
samtools sort SRR7948877.BAM -o sd_2_SRR7948877.BAM
samtools index sd_2_SRR7948877.BAM -o sd_2_SRR7948877.BAM.bai
```

2. Running damidseq_pipeline (Marshall and Brand 2015)

Instructions for using the terminal script damidseq_pipeline can be found here: https://owenjm.github.io/damidseq_pipeline/

The provided GATC fragment file was used in the analysis: D.melanogaster r6 to reflect the most current version of the genome. The Bowtie2 indices required were downloaded from iGenomes (https://support.illumina.com/sequencing/sequencing_software/igenome.html) using the Ensembl BDGP6 version. The fusion BAM and corresponding Dam-only control files were then run using damidseq_pipeline for each replicate. The subsequent software, find_peaks (https://github.com/owenjm/find_peaks) was then used on each of the output files to identify peaks. Genes were identified using the peak file and running the peaks2genes script, using the provided genes list DmR6.genes.gff.gz.

The results from the replicates were compiled using R.

```
Peaks_1 <- rtracklayer::import("path/peaks_1.gff")
```

Peaks_2 <- rtracklayer::import(“path/peaks_2.gff”)

The union of the two GRanges objects was identified with *plyranges::union_ranges()*. This new set of peaks was then compared to the separate replicates and filtered to only contain peaks that overlap with both replicates. The FDR for each peak was selected as the smallest FDR from the two replicates, and the peak’s score was taken from the replicate that matched the FDR value.

The gene lists were similarly filtered to only be the genes in both replicates. The gene lists only contained a gene name and a score. As the scores were not unique in the genes or the peaks, there was no meaningful way to connect the genes and peaks.

3. Running Vissers’ et al., (2018) pipeline

Code used in Vissers’ pipeline can be found here:
https://github.com/jibsch/DamID_Seq_Analysis

As the example analysis begins with count data, the Damsel function *countBamInGatc()* was used to extract counts from the BAM files generated from the processing explained above. The example analysis script was then run. The Python peak caller script was downloaded and used from within the Terminal application. Following the reading of the peak table into R manual adjustment was required to add the chromosome names back into the results. The method used to identify genes (bedtools) proved too unwieldy, so a simplified version of the concept used in Damsel’s *annotatePeaksGenes()* was instead used, by identifying the closest gene for each significant peak using *pair_nearest()* function from the R package plyranges. Damsel’s gene ontology testing was done on this result.

4. Running Damsel

See the Damsel vignette for the code and plots built as a result of the analysis.

For the false discovery rate testing - we switched the sample labels and ran the analysis.

5. Comparison of methods

R code for the comparison of the results between methods is attached at the end of this document.

**SUPPLEMENTARY TABLES:**

Supplementary Table 1. Comparison of functionality between Damsel and previously published methods

| | **Vissers et al., (2018)** | **damidseq_pipeline (Marshall & Brand 2015)** | **Damsel** |
|---|---|---|---|
| Remain within R | ✗ | ✗ | ✓ |
| Generate available counts tables from BAM file | ✗ | ✗ | ✓ |
| Identify regions of enrichment | ✓ | ✓ | ✓ |

| | | | |
|---|---|---|---|
| Incorporate replicates into statistical testing | ✓ | ✗ | ✓ |
| Combine significant regions into peaks | ✓ | ✓ | ✓ |
| Rank peaks based on significance | ✗ | ✗ | ✓ |
| Identify potential gene targets | ✓ | ✓ | ✓ |
| Annotate peaks with gene information | ✗ | ✗ | ✓ |
| Identify enriched gene ontology terms | ✗ | ✗ | ✓ |
| Visualise the results | ✗ | ✗ | ✓ |

**SUPPLEMENTARY FIGURES:**

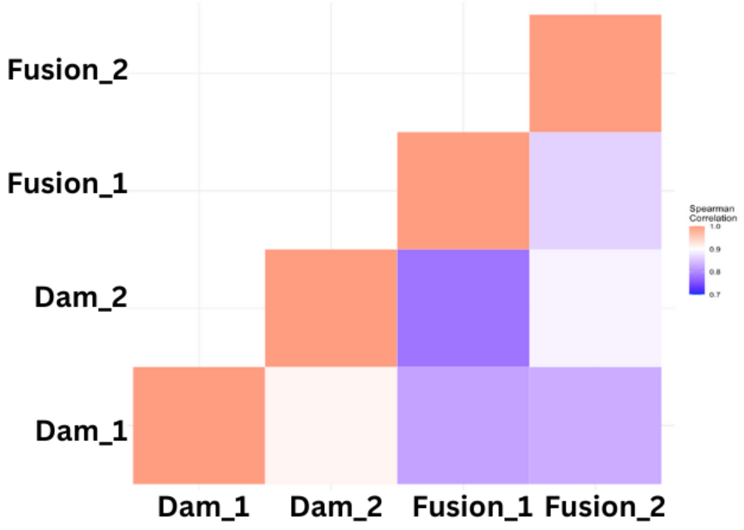


**Supplementary Figure S1.** Heatmap correlation of the region read counts across the genome in Drosophila brain from Vissers et al., (2018). Correlations between all pairs of samples are above 0.7. The Dam_1(control) and Dam_2 (control) samples are very highly correlated (0.9) while the Dam and fusion pairs are slightly less correlated..

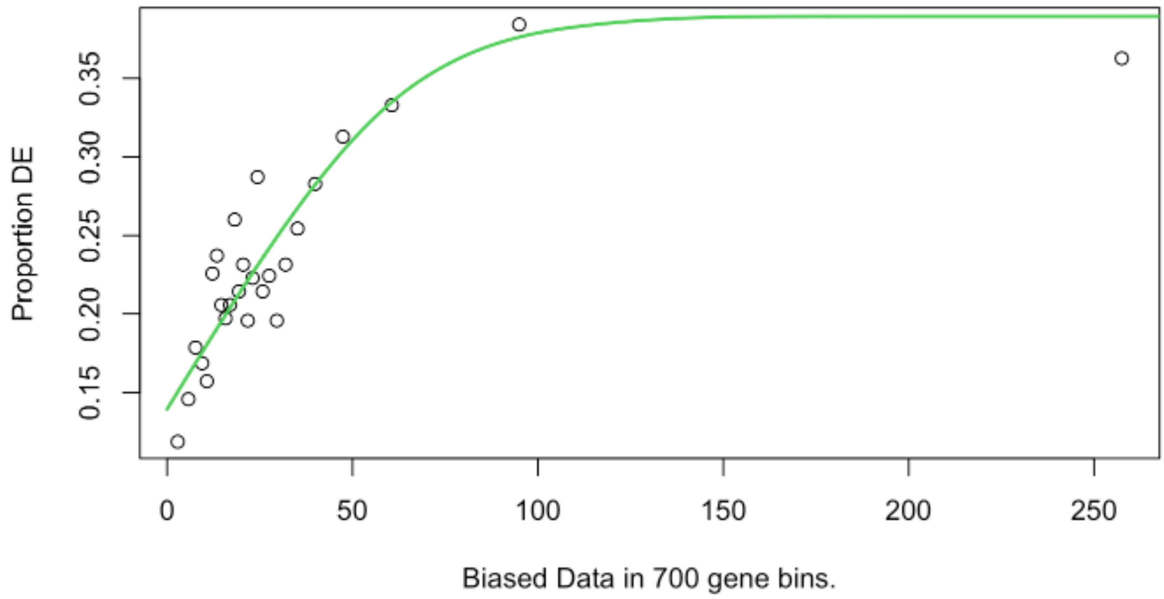


**Supplementary Figure S2. Gene ontology testing**
Plot of the goodness of fit of the model used by goseq. This demonstrates the length based bias present in the data, as genes that contain more GATC regions (x-axis) are more likely to be identified as significant (the proportion on the y-axis). If there was no bias, the line would be horizontal.

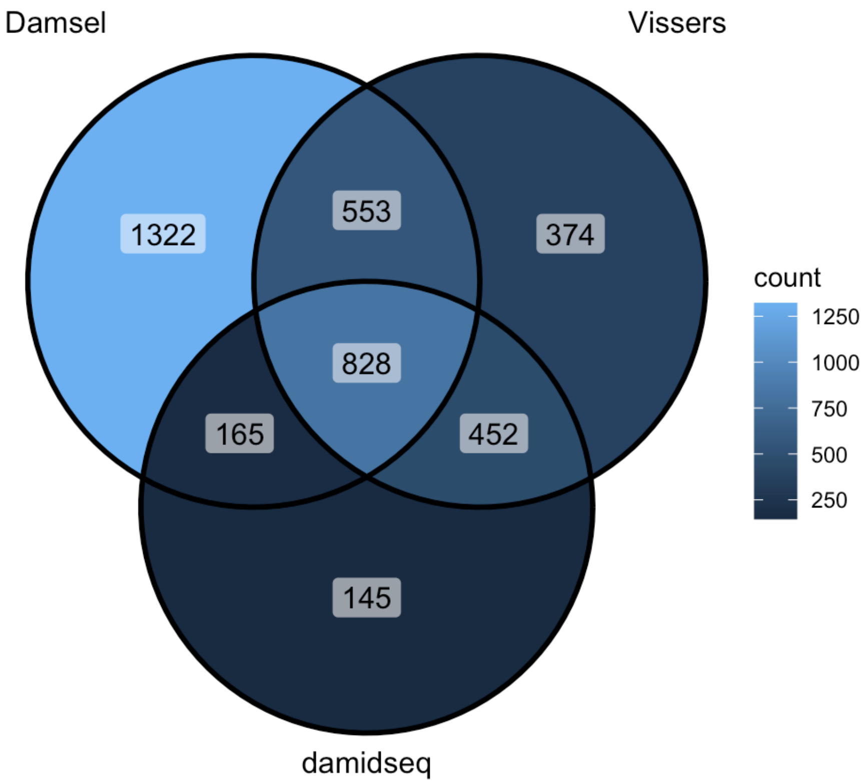


**Supplementary Figure S3. Peak overlap between published methods**
Venn diagram of peak overlap from the analysis of Vissers (2018) data by Damsel, Vissers pipeline, and damidseq_pipeline (Marshall and Brand 2015). As expected, Damsel has a

greater overlap with Vissers as they rely on the same testing procedure (edgeR) than it does with damidseq_pipeline.

**A**

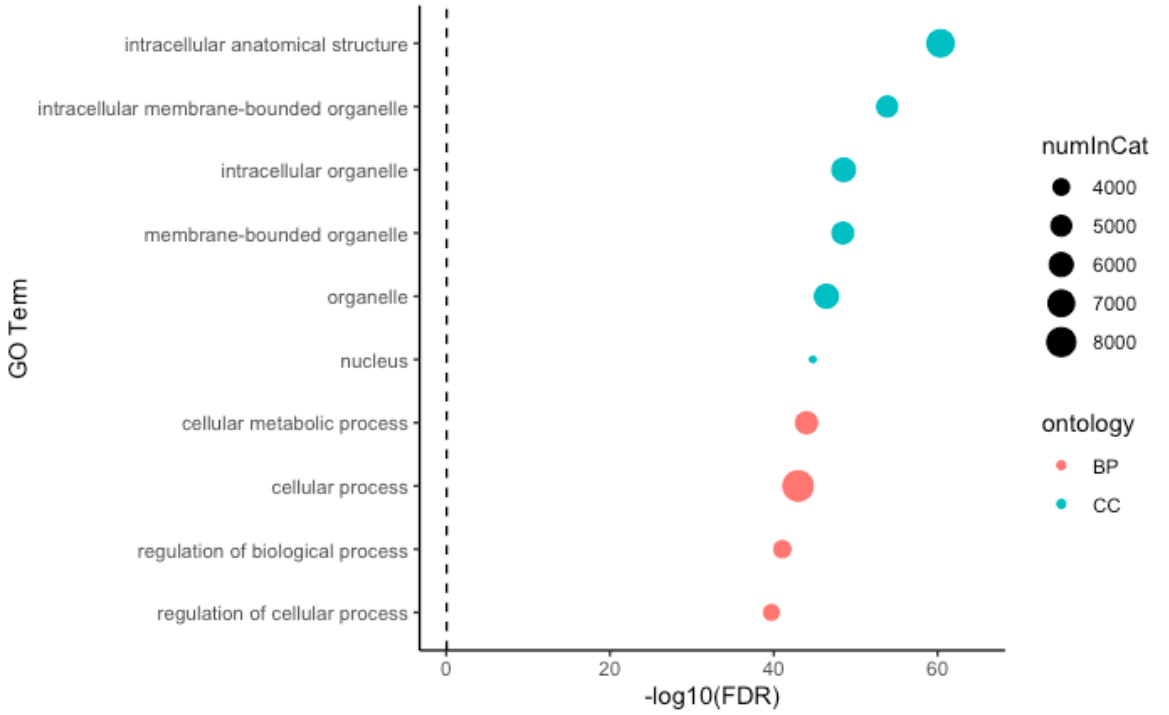


**B**

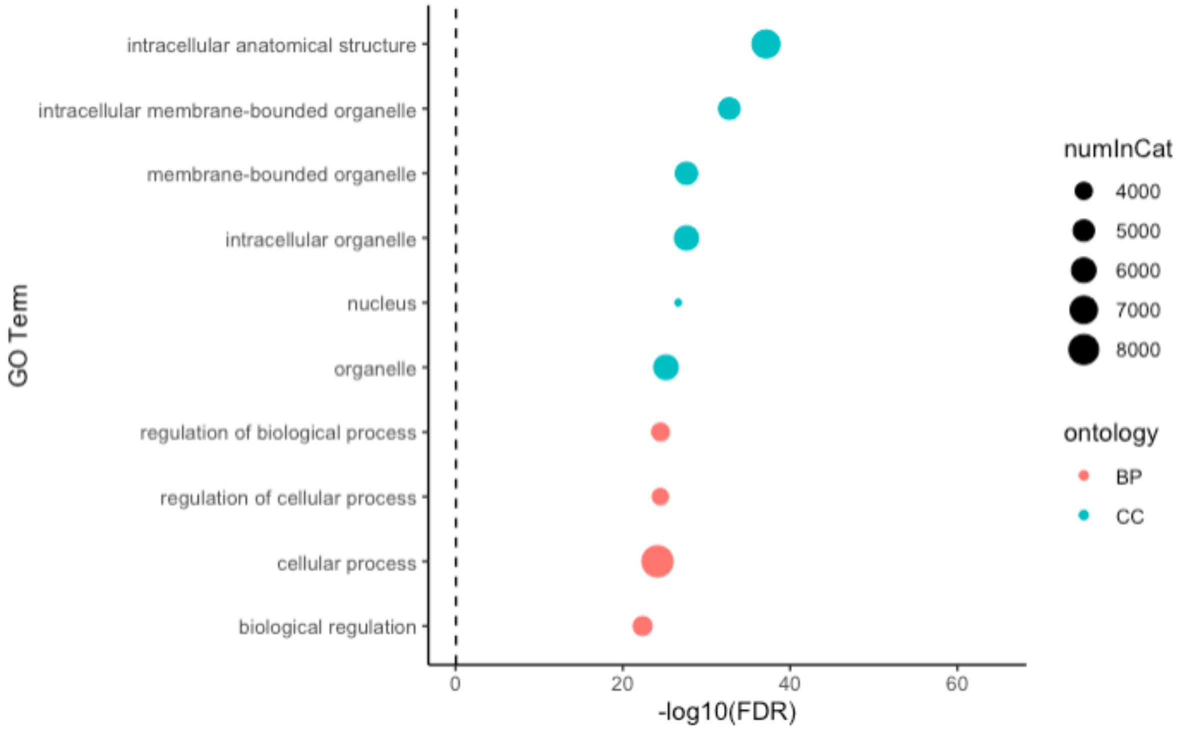


**Supplementary Figure S4.** A) Damsel top 10 gene ontology terms. B) Vissers et al., (2018) top 10 gene ontology terms. The dotted line indicates the FDR value, set to 0.01. Note the overlap of terms between A and B.

### Supplementary Code

````
#########
```{r}
library(plyranges)
library(dplyr)
library(ggVennDiagram)
```

###
# compare peaks
```{r}
bioc_peaks

jan_peaks
jan_peaks_mod

bioc_marshall_1 <- rtracklayer::import("../../find_peaks-master/sd_1_SRR794884-vs-Dam.kde-norm.gatc-FDR0.01.peaks.gff")
bioc_marshall_2 <- rtracklayer::import("../../find_peaks-master/sd_2_SRR7948877-vs-Dam.kde-norm.gatc-FDR0.01.peaks.gff")

bioc_marshall_union <- union_ranges(bioc_marshall_1, bioc_marshall_2) %>% data.frame() %>% mutate(num = 1:n()) %>% as_granges()

bioc_marshall_overlap1 <- pair_overlaps(bioc_marshall_union, bioc_marshall_1) %>% data.frame()
bioc_marshall_overlap2 <- pair_overlaps(bioc_marshall_union, bioc_marshall_2) %>% data.frame()

bioc_marshall_union <- bioc_marshall_union %>% data.frame() %>% mutate(one = bioc_marshall_overlap1[match(.$num, bioc_marshall_overlap1$num), "FDR"],
                                               two = bioc_marshall_overlap2[match(.$num, bioc_marshall_overlap2$num), "FDR"],
                                               one = as.double(one), two = as.double(two)) %>%
  mutate(FDR = case_when(is.na(one) ~ two, is.na(two) ~ one, one <= two ~ one, TRUE ~ two)) %>% replace(is.na(.), 1) %>%
  mutate(source = ifelse(FDR == one, "one", "two")) %>%
  mutate(score = ifelse(source == "one", bioc_marshall_overlap1[match(.$num, bioc_marshall_overlap1$num), "score"], bioc_marshall_overlap2[match(.$num, bioc_marshall_overlap2$num), "score"])) %>%
  mutate(seqnames = paste0("chr", seqnames)) %>% .[,c(1:6,9,11)] %>% .[order(.$FDR),] %>% mutate(rank_fdr = 1:n())
bioc_marshall_union

compare_union <- union_ranges(union_ranges(as_granges(bioc_peaks), as_granges(jan_peaks_mod)), as_granges(bioc_marshall_union)) %>% data.frame() %>%
  mutate(num2 = 1:n()) %>% as_granges()
compare_union

bioc_ov_dam <- pair_overlaps(compare_union, as_granges(bioc_peaks)) %>% data.frame()
bioc_ov_jan <- pair_overlaps(compare_union, as_granges(jan_peaks_mod)) %>% data.frame()
bioc_ov_mars <- pair_overlaps(compare_union, as_granges(bioc_marshall_union)) %>% data.frame()

nrow(jan_peaks_mod)
nrow(bioc_peaks)
nrow(bioc_marshall_union)
2955-2286
828+553
1381/2286
1381/2955
828+165
993/2955
993/1733

venn_data <- list(damsel = bioc_ov_dam$num2, jan = bioc_ov_jan$num2, mars = bioc_ov_mars$num2)
ggVennDiagram(venn_data)

venn_data <- list(Damsel = bioc_ov_dam$num2, Vissers = bioc_ov_jan$num2, damidseq = bioc_ov_mars$num2)
ggVennDiagram(venn_data, label = "count")
```
# compare genes
```{r}
bioc_mars_genes_1 <- read.table("../../find_peaks-master/sd_1_SRR794884-vs-Dam.kde-norm.gatc-FDR0.01.peaks.genes.in.peak.csv")

bioc_mars_genes_2 <- read.table("../../find_peaks-master/sd_2_SRR7948877-vs-Dam.kde-norm.gatc-FDR0.01.peaks.genes.in.peak.csv")

venn_data <- list(damsel = bioc_anno$all$gene_name, jan = jan_peaks_anno$all$gene_name, mars = rbind(bioc_mars_genes_1, bioc_mars_genes_2)$V1)
ggVennDiagram(venn_data)

venn_data <- list(damsel = bioc_anno$closest$gene_name, jan = jan_peaks_anno$closest$gene_name, mars = rbind(bioc_mars_genes_1, bioc_mars_genes_2)$V1)
ggVennDiagram(venn_data)

venn_data <- list(damsel = bioc_anno$all$gene_name, jan = jan_peaks_anno$closest$gene_name, mars = rbind(bioc_mars_genes_1, bioc_mars_genes_2)$V1)
ggVennDiagram(venn_data)

length(unique(bioc_anno$closest$gene_name))
length(unique(jan_peaks_anno$closest$gene_name))
length(unique(rbind(bioc_mars_genes_1, bioc_mars_genes_2)$V1))
(419+503)/2081
(419+503)/1700
(419+182)/2081
(419+182)/2704

venn_data <- list(damsel = filter(bioc_ov_dam, rank_p <= 500)$num2, jan = filter(bioc_ov_jan, rank_p <= 500)$num2, mars = filter(bioc_ov_mars, rank_fdr <= 500)$num2)
ggVennDiagram(venn_data)

venn_data <- list(damsel = filter(bioc_ov_dam, rank_p <= 1000)$num2, jan = filter(bioc_ov_jan, rank_p <= 1000)$num2, mars = filter(bioc_ov_mars, rank_fdr <= 1000)$num2)
ggVennDiagram(venn_data)
```
````
